# Experimental evidence for the role of natural radioactivity in influencing viability of commensal microorganisms

**DOI:** 10.1101/2020.07.21.214676

**Authors:** Marco Ruggiero

**Author notes:** **Corresponding author**: Marco Ruggiero, MD, PhD., National Coalition of Independent Scholars, San Antonio, TX, USA.

## Abstract

The role of natural radioactivity in influencing viability of commensal microorganisms, with particular reference to those pertaining to the human microbiota, has not been investigated in detail. The results of experiments culturing a diversified array of microorganims - probiotics - in naturally radioactive mineral water or deuterium-depleted water are here described. Culturing microorganisms in naturally radioactive mineral water yielded one order of magnitude more live microbial cells in comparison with culturing in deuterium-depleted water. Based on these experimental results, a method for co-culturing prebiotic microorganisms (cyanobacteria) that are extremely resistant to the harmful effects of radiations, together with the probiotics mentioned above, is described. The goal of this co-culture in naturally radioactive mineral water is to transfer the information from the radiation-resistant microorganisms to the probiotics whose viability is enhanced by natural radioactivity. Expression of DNA repair genes in cyanobacteria is induced by co-culturing in a medium of carbonated mineral water naturally containing the radioactive isotopes ^228^U and ^226^Ra. The culture medium, in addition to naturally radioactive water, contains silica from vegetal origin to enhance horizontal gene transfer. Finally, it is described the transfer of resistance to the harmful effects of radiations to human cells through a Lactococcus phage-encoded protein, ORF252, that is the evolutionary precursor of human proteins involved in DNA repair and cancer protection.

## Introduction

Static magnetic fields, ionizing, and non-ionizing radiations exert biological effects on normal and transformed cells in culture, embryos and organisms, inducing changes in signal transduction and gene expression (1–24). Certain types of eukaryotic cells, notably certain types of cancer cells (3–11), show resistance to the killing effects of ionizing radiations, a phenomenon that, in the context of radiotherapy of cancer, may represent an obstacle to effective treatment (14). Most prokaryotic cells are sensitive to the killing effects of ionizing radiations and even to weak electromagnetic fields such as those generated by the physiologic electric activity of human cells *in vivo* (19). However, certain prokaryotes such as cyanobacteria, exhibit extremely high resistance to the killing effects of ionizing radiations (25). A notable example is represented by Arthrospira platensis, a bacterium that survives exposure to gamma-rays up to 6,400 Gy with a dose rate of 527 Gy h^−1^ (25) [for reference, the median human lethal radiation dose computed from data on occupants of reinforced concrete structures in Nagasaki, Japan, is around 3 Gy (26)]. Investigation in the behavior of commensal microorganisms that can be defined probiotics challenged with naturally radioactive mineral water was prompted by the ample spectrum of responses to radiations exhibited by microorganisms. Probiotics are consumed in increasingly large amount for their health supporting properties (27), and since their efficacy lays in the ability to colonize the intestine, it is interesting to assess whether drinkable radioactive water affects their viability. In order to assess the role of radioactivity in the responses of commensal microorganisms, the results obtained culturing an array of microorganisms in radioactive mineral water were compared with the results obtained culturing the same array of microorganisms in deuterium-depleted water. Furthermore, the role of Arthrospira platensis in conferring radiation resistance was evaluated. This bacterium, in addition of being extraordinarily resistant to ionizing radiations, is considered a prebiotic since it favors the growth of probiotic bacteria. Therefore, it can be hypothesized that co-culturing Arthrospira platensis with probiotics, in the presence of radioactive mineral water, may yield information on possible horizontal transfer of information between radio-resistant and radio-sensitive microorganisms.

## Materials and Methods

### Radioactive mineral waters

Two types of radioactive mineral waters were used. Analyses for minerals and radioactivity were performed by the Institut für Pflanzenernährung und Bodenkunde, Braunschweig, Germany (28). The first type of radioactive water, termed “A”, was used to compare the viability of cells of an array of microorganisms cultured in radioactive water *versus* the same microorganisms cultured in deuterium-depleted water. Composition of the first water, termed A, as far as minerals and radioactive isotopes are concerned was the following:

|  |  |
| --- | --- |
| pH | 5.5 |
| HCO<sup>3+</sup> | 483 mg/L |
| Ca<sup>2+</sup> | 486 mg/L |
| Cl<sup>-</sup> | 8.6 mg/L |
| Mg<sup>2+</sup> | 84 mg/L |
| Na<sup>+</sup> | 9.1 mg/L |
| K<sup>+</sup> | 3.2 mg/L |
| SO<sup>4-</sup> | 1,187 mg/L |
| <sup>226</sup>Ra | 5.5 mBq l<sup>-1</sup> |
| Uranium | 1.40 mBq l<sup>-1</sup> |
| Main minerals | 2,181 mg/L |
| Radioactivity from <sup>226</sup>Ra + Uranium (<sup>238</sup>U): | 40.9 mBq l<sup>-1</sup> |
| Radioactivity from Uranium: | 86% |

The second type of radioactive water, termed B, was used for co-culturing Arthrospira platensis with a blend of probiotic microorganisms whose composition is described below. Composition of the second water, termed B, as far as minerals and radioactive isotopes are concerned was the following:

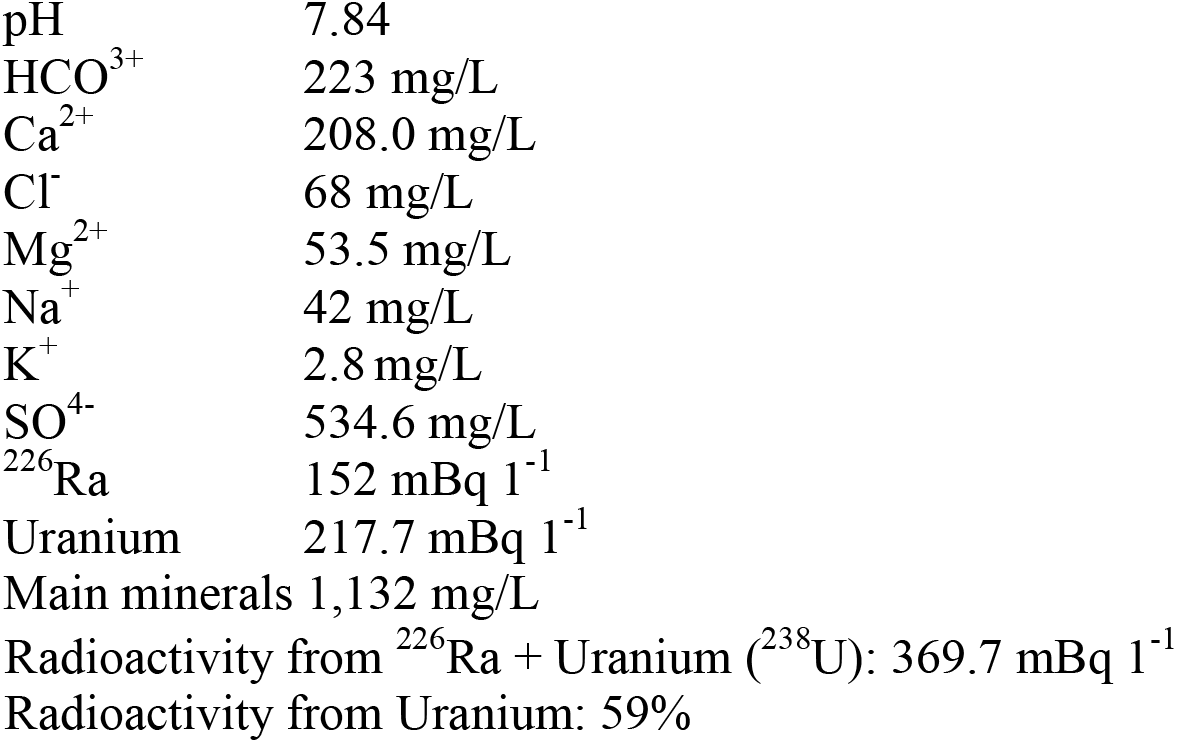

At variance with water A, water B was carbonated. The rationale to choose a carbonated mineral water with these values of natural radioactivity lays in the goal of stimulating adaptogenic responses in radio-resistant cyanobacteria that are known to up-regulate genes involved with DNA repair and protection as consequence of exposure to radiations (29).

### Deuterium-depleted water

Deuterium-depleted water was purchased from Litewater Scientific (Reno, NV, USA). According to the Company’s website (www.drinklitewater.com) this water has a 94-97% lower concentration of deuterium than regular water. This water is obtained from a natural artesian well in Russia, in the state of Tambov; it is purified by membrane technology and then processed by a method known as low-temperature vacuum rectification that is claimed to remove up to 97% of the deuterium in water.

### Characteristics of the probiotic blend used for co-culturing with Arthrospira platensis

The probiotic blend was kindly donated by Silver Spring Sagl (Arzo-Mendrisio, Switzerland); it contains microorganisms that are members of the families of Streptococcaceae, Lactobacillaceae, Leuconostocaceae, Bifidobacteriaceae, and Thermaceae. The blend contains also lyophilized kefir grains that are characterized by a complex composition. The unique microbial composition of this probiotic blend was fully described in two recent articles (30,31). Human consumption of this blend is associated with detoxification of non-metal toxicants (32), decrease of serum alpha-N-acetylgalactosaminidase (nagalase) activity (32,33), decrease of serum C-reactive protein (CRP) (33), decrease of markers specific for multiple myeloma (34), and decrease of viral load in Hepatitis B (35).

### Determination of microbial viability in radioactive water A, and deuterium-depleted water

Microbial cultures, defined as culture starter, were kindly donated by Silver Spring Sagl (Arzo-Mendrisio, Switzerland). In one arm of the experiment, radioactive water A was used; in the other arm, deuterium-depleted water was used. 45 g of hemp seed proteins were added to 500 mL of each type water. The mixture of water and hemp seed proteins was boiled for 10 s; after that, it was cooled at room temperature (22 °C). When the mixture had reached room temperature, the following ingredients were added: lemon juice (7.5 g); cane sugar (25 g); apple cider vinegar (5 g); culture starter (a blend of active cultures, cultured kefir grains with Lactobacillus bulgaricus, Streptococcus thermophilus, Bifidobacterium infantis, Bifidobacterium bifidum, Bifidobacterium lactis, Bifidobacterium longum) (2.5 g). The mixture was left at room temperature for a total of 48 hours; after the first 24 h, the mixture was gently stirred. After that, the fermented product was gently mixed and then lyophilized using an apparatus provided by Zhengzhou Hento Machinery Ltd (Zhengzhou City, Henan, China). Yield was 12.64 % for the preparation using radioactive water A, and 12.69% for the preparation using deuterium-depleted water. The lyophilized powders, one containing the microbes cultured in radioactive water A, and the other the microbes cultured in deuterium-depleted water, were then analyzed by flow cytometry for the count of viable cells (Biolab Research Srl, Novara, Italy).

### Method for co-culturing in radioactive water B

The method described here is for 1 L of product. In case of hypothetical future industrial manufacture, the procedure could be scaled up indefinitely. As first step, the radioactive carbonated mineral water designated B is heated up to 39°C. Then, the following ingredients are added: 40 g of Equisetum arvense; 40 g of white sugar; 15 mL of lemon juice; 10 mL of apple cider vinegar; 45 g of Arthrospira platensis; 10 mg of the probiotic blend described above. Once these ingredients have been added, the solution is gently mixed by stirring at time 0, that is immediately after addition of the ingredients, 12, 24, and 36 h thereafter. Temperature of the mixture is maintained between 36°C and 39°C for a total incubation time of 48 h. After 48 h incubation, the solution is gently mixed and lyophilized using the method described above.

### Study of amino acid sequences and protein structures

Study and comparisons of amino acid sequences, alignments and study of protein structures were performed using the dedicated programs developed by The Universal Protein Resource (UniProt) Consortium, a comprehensive resource for protein sequence and annotation data (36).

## Results

Fig. 1, shows the products obtained by culturing microorganisms in radioactive mineral water (A) and deuterium-depleted water, as they appear after lyophilization.

**Fig. 1.**
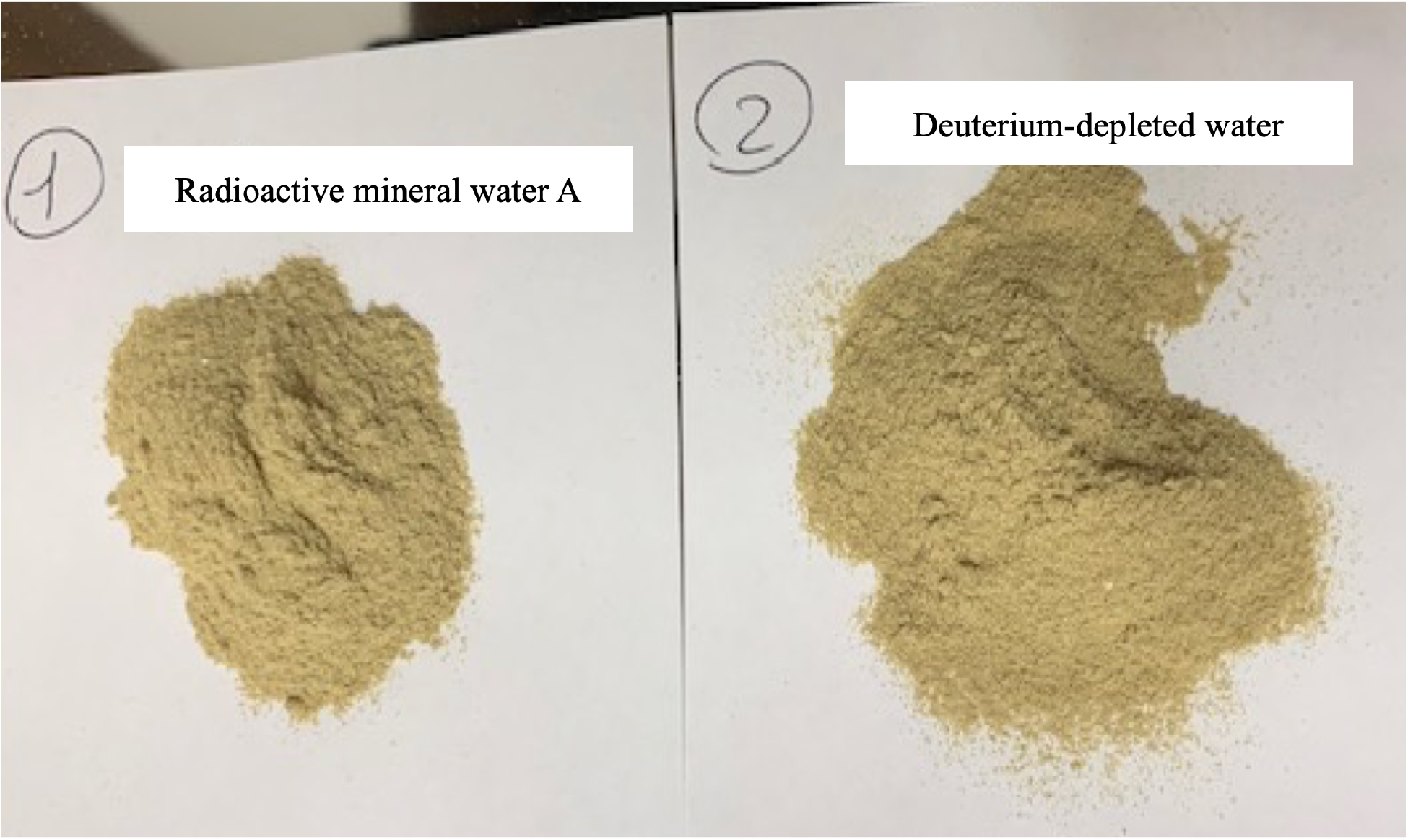
Appearance of the products of lyophilization of microorganisms cultures in radioactive mineral water designated A (left panel) and deuterium-depleted water (right panel). Culture conditions and procedure of lyophilization are described in the Materials and Methods. The two powders appear identical.

Determination of viable cells by flow cytometry demonstrated that in the sample derived from culturing in radioactive mineral water A there were 2.36 × 10^9^ viable cells per g of lyophilized powder; in the sample derived from culturing in deuterium-depleted water, there were 8.48 × 10^8^ viable cells per g of lyophilized powder. These results demonstrate that radioactive mineral water significantly favors viability of commensal microorganisms.

Based on these results, experiments were performed to assess the feasibility of co-culturing the probiotic blend described in references 30-35 together with Arthrospira platensis using a much more radioactive water, that is the water designated B, whose composition is described in Materials and Methods. The rational for such an experiment lays in the continuous and simultaneous exposure to electromagnetic radiations (from radioactivity), and induction of photosynthesis from the Cherenkov radiation that is also due to natural radioactivity. Thus, in the absence of Cherenkov radiation, cyanobacteria such as Arhtrospira down-regulate the genes presiding to photosynthesis when exposed to radiations such as gamma-rays and, at the same time, up-regulate the genes that protect them from the harmful effects of radiations (29). In the experiment described here, however, induction of photosynthesis occurs concomitantly with exposure to radiations since induction of photosynthesis is a by-product of superluminal charged particles emitted by the radioisotopes of the water. It is well known that the process of photosynthesis is based on quantum entanglement and, therefore, it can be hypothesized that up-regulation of genes involved in DNA repair and resistance to radiations, and induction of photosynthesis become entangled. This may lead to further entanglement between the radio-resistant cyanobacteria and the microorganisms of the probiotic blend that are also exposed to the radioisotopes of water but have no photosynthetic capacity; the mechanism at the basis of living bacterial entanglement following exposure to electromagnetic fields is explained in detail in Marletto *et al.* (37). The choice of naturally radioactive water that is also carbonated lays in the well-known concept that carbon dioxide is required for photosynthesis and such a requirement is all the more significant when the source of light is represented by the continuum spectrum of Cherenkov radiation. In addition, carbonated water shows a distinctive light scattering pattern that is very different from that of non-carbonated water. The presence of dissolved gas in the water leads to smoothing of the angular dependence of scattered light intensity, and reduction of the polydispersity degree, thus optimizing the conditions for Cherenkov radiation-induced photosynthesis (38). Such an optimization is due to the uniform distribution in the water of the radioactive isotopes of Uranium and Radium as well as to the equally uniform distribution of the bubbles typical of carbonated water. The constant, uniform, Cherenkov radiation is scattered by the bubbles and therefore reaches the light harvesting antennae of the cyanobacteria from all possible angles at variance with a common source of light that would reach the antennae from a fixed angle.

In short, the natural radioactivity in the carbonated mineral water evokes radio-protective responses as well as induction of photosynthesis in the cyanobacteria; through quantum entanglement, these transfer the radio-protective response to the microorganisms of the probiotic blend that are also exposed to the natural radioactivity but are not radio-resistant *per se*.

As far as the use of Arthrospira platensis is concerned, it is worth noting that it has been used as food since the 16^th^ century, because of its high-protein content and digestibility, and it is considered a prebiotic because it favors the growth of known probiotic microorganisms such as lactic acid bacteria [Lactococcus lactis, Streptococcus thermophilus, Lactobacillus casei, Lactobacillus acidophilus, and Lactobacillus bulgaricus (39)]. All these microorganisms are part of the probiotic blend described in references 30-35, and are co-cultured with Arthrospira in the naturally radioactive carbonated mineral water designated B. Consistent with this approach, Arthrospira platensis has gained increasing interest both for the general public and, more specifically, for astronauts who may be exposed to electromagnetic radiations (40). In specific regard to protection against the harmful effects of electromagnetic radiations, its strong anti-oxidant and anti-inflammatory properties show potential application in human radiation protection (41). Resistance to radiations is due to a number of factors that include the presence of high level of anti-oxidants and, more importantly, induction and up-regulation of genes involved in DNA repair. Among these, of particular importance in the context of this study are the genes of the *uvr* family that are involved in repair of single strand breaks (SSBs) of DNA. DNA SSBs are the most typical DNA lesions observed after exposure to many types of electromagnetic radiations including those of mobile telecommunication (42), and it is postulated that the risks for human health due to exposure to such radiations derive from this type of genotoxic effects. To this point, the International Agency for Research on Cancer (IARC), after having evaluated the available scientific data, has classified these electromagnetic fields as being “possibly carcinogenic” to human (43). These effects are of particular interest in the context of the fifth generation technology standard for cellular networks, which cellular phone companies began deploying worldwide in 2019. Although no studies are available on the direct effects of this new technology standard on human cancer onset and progression, a recent peer-reviewed article analyzed 94 relevant publications performing *in vivo* or *in vitro* research on this topic. According to this meta-analysis, 80% of the *in vivo* studies showed biological responses to exposure, while 58% of the *in vitro* studies demonstrated effects. All the responses showed effects on all biological endpoints analyzed (44). In addition, a study published in May 2020 in the peer-reviewed journal “Toxicology Letters” by Authors from the Georgia Institute of Technology of the United States, McGill University of Canada, Albert Einstein College of Medicine of the United States, University of Crete of Greece, and Sechenov University of Moscow, Russia, states that “*… the nascent mobile networking technology will affect not only the skin and eyes, as commonly believed, but will have adverse systemic effects as well*” (45).

The rationale for the use of Arthorspira platensis in co-culture with the probiotic blend consists in the following considerations. Ionizing radiations as well as electromagnetic radiations used for mobile telecommunication and, in particular, high frequency fields used for the fifth generation technology standard, affect biological responses and are classified as carcinogenic for humans (43). The most common types of DNA damage due to electromagnetic radiations used for mobile telecommunication are SSBs (42). This damage is repaired by proteins encoded by *uvr* genes and Arthrospira platensis up-regulates expression of *uvr* genes upon exposure to radiations (29). Arthrospira platensis is known to protect against radiation sickness (46) as well as against the damage inflicted by electromagnetic radiations through DNA repair (*uvr* gene up-regulation) (47). Naturally radioactive carbonated mineral induces up-regulation of radiation protection genes in Arthrospira platensis and Cherenkov radiation induced photosynthesis (48). Photosynthesis is based on quantum entanglement (49). Electromagnetic fields such as those associated with charged particles from radioisotopes as those in the mineral water induce quantum entanglement between microorganisms (37). Quantum entanglement and transfer of resistance to radiation between living organisms occurs in water (50). Therefore, Arthrospira platensis, a prebiotic bacterium already known to protect from radiation sickness, once exposed to (that is cultured in) naturally radioactive mineral water, activates mechanisms for DNA protection (*uvr* genes) that are known to protect against SSBs, and transfers such a protection to probiotic microorganisms co-cultured in the same medium that in turn transfer such a protection to human cells through phages as shown below.

The rationale for the use of the probiotic blend described in references 30-35, in co-culture with Arthrospira platensis consists in the following considerations. The human microbiota is negatively affected by electromagnetic radiations of all types including those of mobile telecommunication (51). The probiotic blend described in references 30-35 contains a high number of live probiotic microorganisms with a high degree of biodiversity. Such a biodiversity reproduces the complexity of the human microbiota, thus reconstituting an organ that is essential for human health (52). The probiotic blend contains plasmids that are known to transfer biological information between bacteria (30), and this feature is essential for exchanging biological information pertaining to radiation protection. In addition, the probiotic blend contains phages that are known to stimulate immunity, repair DNA and transfer biological information to human cells (31). Independent of the route of administration, phages enter the bloodstream and tissues and interact with immune cells (macrophages, T-cells, Natural Killer cells, B-cells, granulocytes) (53). Phages have anti-inflammatory properties and inhibit excessive reactive oxygen species production (54), and it is known that excessive reactive oxygen species production is one of the consequences of exposure to electromagnetic radiations and is responsible for damage to DNA, proteins and cell structures (55,56). In addition, the probiotic blend described in references 30-35, contains naturally formed Gc protein-derived Macrophage Activating Factor, a cytokine that stimulates/rebalances the immune system and in the probiotic blend is 100 fold more active than the chemically-obtained counterpart (33). Optimization of macrophage function is of utmost importance in counteracting the harmful effects of electromagnetic radiations, including those used in mobile telecommunication, since exposure to these electromagnetic radiations provokes macrophage dysfunction (57) that contributes to immune system disturbances (58) that, in turn, contribute to the adverse health effects of mobile networking technology under real-life conditions (45). Ionizing radiations as well as electromagnetic radiations used for mobile telecommunication cause excessive formation of reactive oxygen species that leads to apoptosis of human blood macrophages (56). The simultaneous presence of phages, that inhibit excessive reactive oxygen species (54), and Gc protein-derived Macrophage Activating Factor that stimulates human blood macrophages (59), prevents these harmful effects of electromagnetic radiations on macrophages. The simultaneous presence of phages and Gc protein-derived Macrophage Activating Factor leads to optimization of immune system function at the same time preventing inflammation (54), a threatening pathologic feature that accompanies dysregulated immune responses to ionizing radiation as well as to mobile telecommunication electromagnetic radiations (60). Co-culturing Arthrospira platensis with the probiotic blend in the same aqueous medium allows exchange of biological information through horizontal gene transfer and quantum entanglement as specifically demonstrated for protective responses to radiations (50). The transfers of information described above are based on the homologies of amino acid sequence and function of the genes belonging to the family of *uvr*. Here, it is demonstrated that proteins coded for by *uvr* genes of Arthospira platensis and Lactococcus lactis subsp. lactis, one of the strains of the probiotic blend described in references 30-35, share significant homologies that are responsible for the transfer of information as shown in the following figures.

Fig. 2, shows the high degree of similarity between UvrABC system protein B (uvrB gene) of Arthrospira platensis (upper line), and UvrABC system protein B (uvrB gene) of Lactococcus lactis subsp. lactis (lower line).

**Fig. 2.**
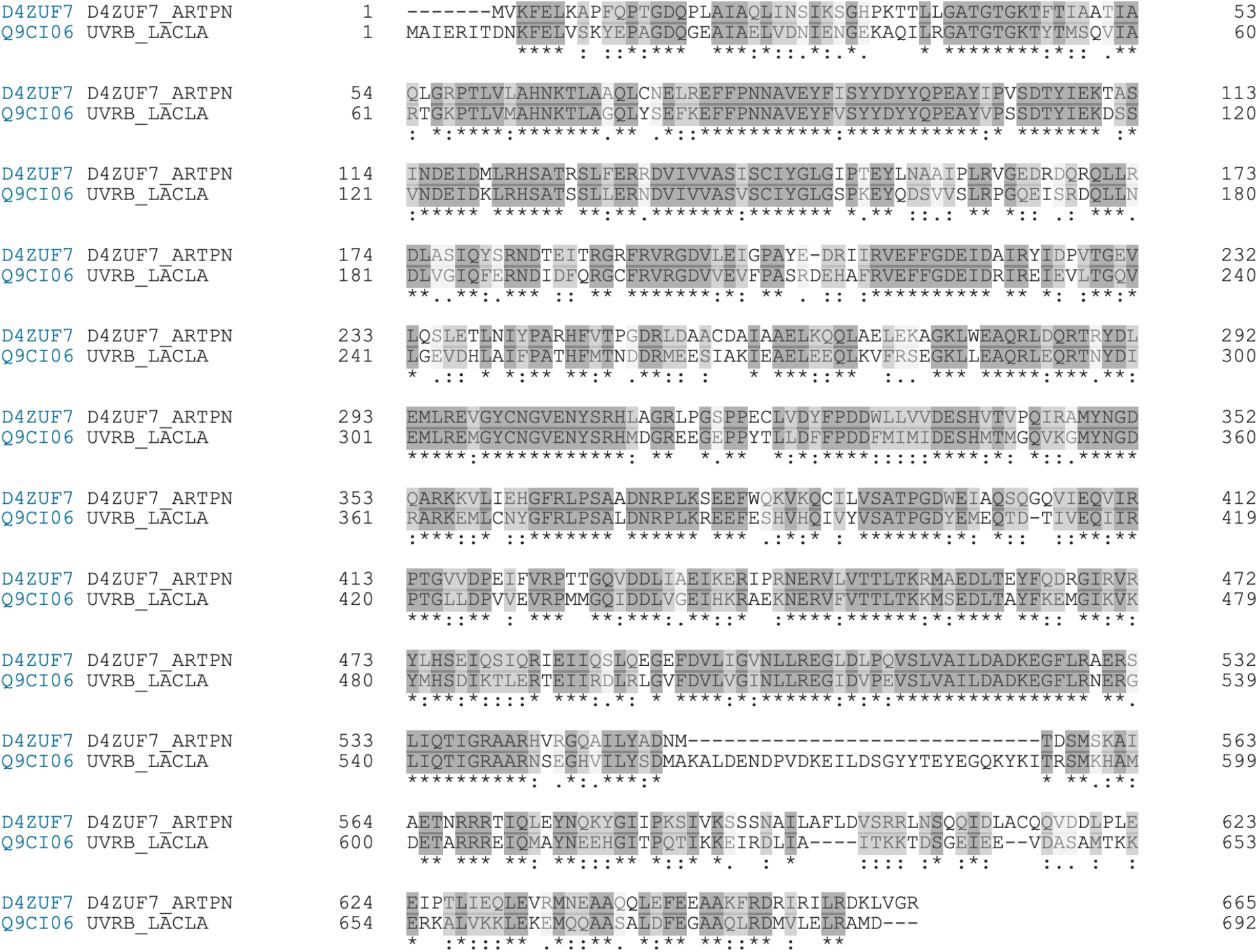
Results of alignment between UvrABC system protein B (uvrB gene) of Arthrospira platensis (upper line), and UvrABC system protein B (uvrB gene) of Lactococcus lactis subsp. lactis (lower line). Residues (amino acids) showing significant degree of similarity are highlighted in grey. Alignment displays the following symbols denoting the degree of conservation observed in each column:

An * (asterisk) indicates positions which have a single, fully conserved residue.
A : (colon) indicates conservation between groups of strongly similar properties - scoring > 0.5 in the Gonnet PAM 250 matrix.
A . (period) indicates conservation between groups of weakly similar properties - scoring =< 0.5 in the Gonnet PAM 250 matrix.

Fig. 3, shows the high degree of homology as far as hydrophobic, positively and negatively charged amino acids are concerned.

**Fig. 3.**
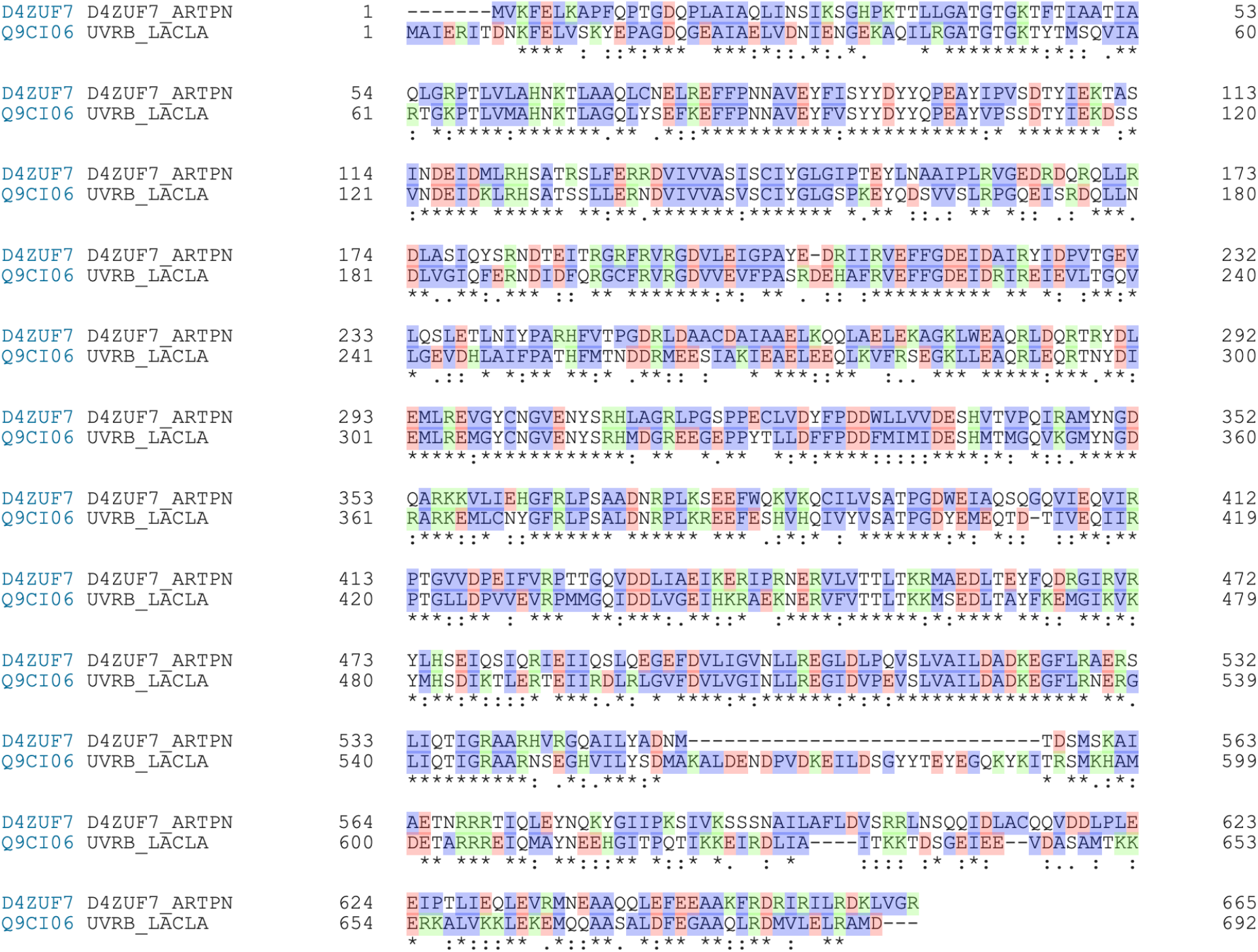
Results of alignment between UvrABC system protein B (uvrB gene) of Arthrospira platensis (upper line) and UvrABC system protein B (uvrB gene) of Lactococcus lactis subsp. lactis (lower line). Hydrophobic residues are highlighted in indigo. Positively charged residues are highlighted in green. Negatively charged residues are highlighted in pink.

Fig. 4, shows the identity of the nucleotide binding domains that are essential for DNA repair.

**Fig. 4.**
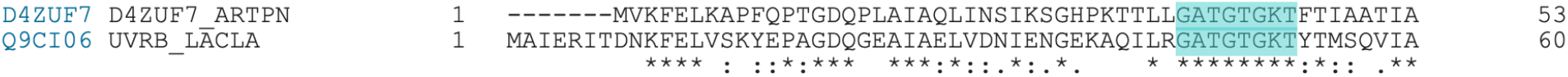
Results of alignment between UvrABC system protein B (uvrB gene) of Arthrospira platensis (upper line) and UvrABC system protein B (uvrB gene) of Lactococcus lactis subsp. lactis (lower line). Nucleotide binding residues are highlighted in cyan.

Fig. 5, shows the high degree of homology in the distribution of big and small amino acids, a feature that determines the spatial configuration of the protein and its ability to bind and repair DNA.

**Fig. 5.**
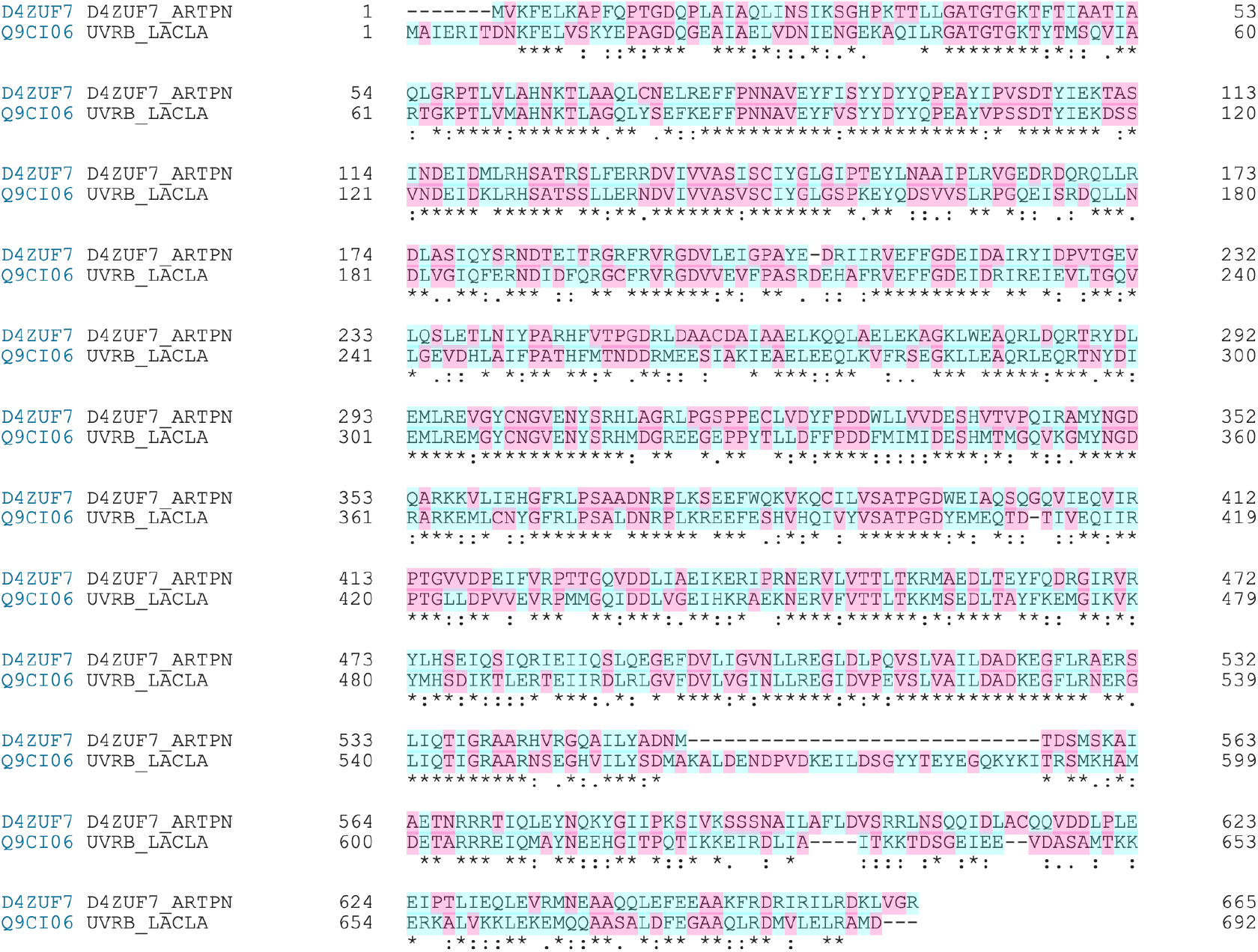
Results of alignment between UvrABC system protein B (uvrB gene) of Arthrospira platensis (upper line) and UvrABC system protein B (uvrB gene) of Lactococcus lactis subsp. lactis (lower line). Small residues are highlighted in pink. Big residues are highlighted in light blue.

Fig. 6, shows the high degree of homology in the motifs of both proteins.

**Fig. 6.**
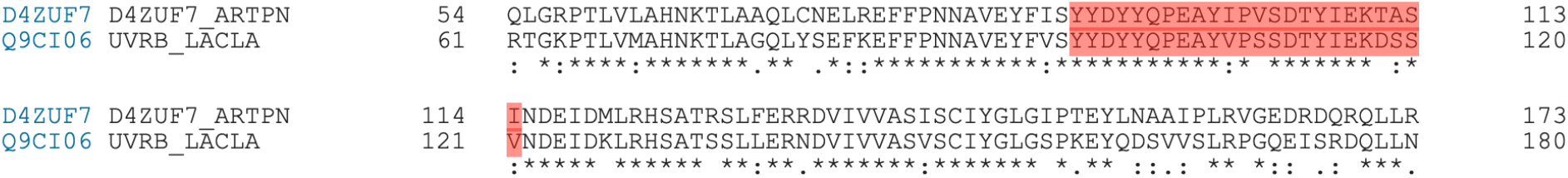
Results of alignment between UvrABC system protein B (uvrB gene) of Arthrospira platensis (upper line) and UvrABC system protein B (uvrB gene) of Lactococcus lactis subsp. lactis (lower line). Motifs are highlighted in red.

The similarities and homologies shown in Figs. 2–6 explain the transfer of information between Arthrospira platensis and one of the constituents of the probiotic blend. The following observations explain the transfer of information to human cells through the intermediary that is constituted by Lactococcus phage ul36.

The complete genomic sequence of the Lactococcus lactis phage ul36 belonging to P335 lactococcal phage species was determined and analyzed in 2002 (61). The genomic sequence of this lactococcal phage contained 36,798 base pairs with an overall G+C content of 35.8 mol %. Fifty-nine open reading frames (ORFs) of more than 40 codons were found. N-terminal sequencing of phage structural proteins as well as bio-informatic analysis led to the attribution of a function to 24 of these ORFs (41%). Among these, of particular importance is the ORF designated ORF252 (Q9MC33). The amino acid sequence of this protein is reported in Fig. 7.

**Fig. 7.**
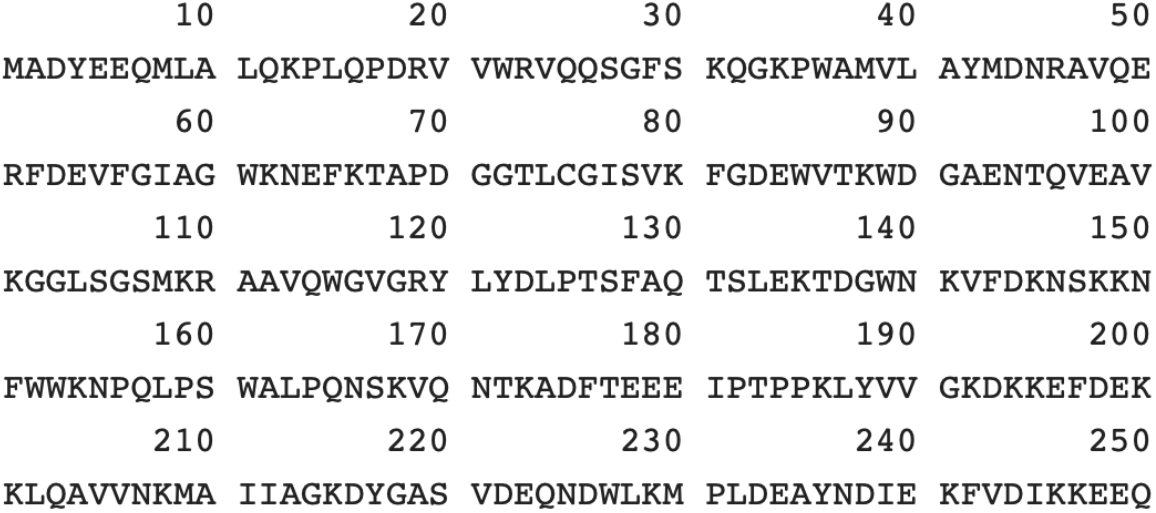
Amino acid sequence of ORF252 of Lactococcus phage ul36.

This protein is a viral ancestor of the human protein RAD52 that is essential for DNA repair, resistance to the harmful effects of electromagnetic radiations, genomic stability and, cancer prevention (62). Purified ORF252 binds single-stranded DNA preferentially over double-stranded DNA and promotes the renaturation of long complementary single-stranded DNAs in the context of DNA repair. Interestingly, ORF252 is able to self-assemble even in the absence of DNA with the formation of toroidal structures (62). This feature of ORF252 is interesting in the context of the interactions described here, with particular reference to DNA repair, in particular, if it is compared to the self-assembly of a major protein contributing to human genome stability, the tumor suppressor protein p53, “*the guardian of the - human - genome*” (63). It was demonstrated that four p53 molecules need DNA to self-assemble on two half-sites of the polynucleotide to form a tetramer that consists of a dimer of dimers whose stabilization is assured by protein-protein and DNA base-stacking interactions (64). It appears that the ability of ORF252 to protect genome stability is more “*primordial*” as the information for self-assembly is all contained in the protein structure without the need for interactions with damaged DNA. This feature provides ORF252 with the advantage of being less restricted in its ability to repair DNA and guarantee genome stability. Self-assembly of ORF252 in toroidal structures raises the interesting possibility of non-chemical signaling between the protein and DNA, possibly through biological quantum entanglement between the electron clouds of toroids of DNA and the electron clouds on the surface of toroidal ORF252 (65). Toroidal structures of DNA and proteins such as ORF252 become entangled because of their spatial geometry; this phenomenon is not dissimilar from the quantum entanglement at the level of the protein tubulin in neuron microtubules, a phenomenon that is thought to be at the basis of human consciousness (66). It is also consistent with a recent paper entitled “*Is the Fabric of Reality Guided by a Semi-Harmonic, Toroidal Background Field?*” (67). It is interesting to note that DNA itself is endowed with an intrinsic degree of consciousness (68) that obviously does not rely upon quantum entanglement at the level of tubulin. It is also interesting to note that ancient viruses may have been responsible for the onset of consciousness in humans (69) and, therefore, it is plausible that the intrinsic consciousness of the DNA of viruses and humans may have become entangled at different levels. As far as the role of ORF252 in the context of DNA repair is concerned, it is worth noting that ORF252 binds RecA and stimulates homologous recombination reactions. Not surprisingly, human RAD52, an evolutionary derivative of ORF252, was recently defined “*a novel player in DNA repair in cancer and immunodeficiency*” and was attributed a tumor suppressor function (70). These functions of ORF252/RAD52 are consistent with a recent clinical observation in a case of multiple myeloma (34). It is also worth noting that RAD52 was suppresses HIV-1 infection in a manner independent of its ability to repair DNA double strand breaks through homologous recombination (31, 71). Although there are no direct evidences for a role of ORF252 protein in suppressing retroviral infections, since this activity resides in the DNA-binding and self-interaction domains of RAD52 that are homologous to ORF252, it is tempting to speculate that ORF252 of phage ul36 may also have anti-retroviral activities by binding to retroviral cDNA and suppressing integration of the infecting virus.

Here it is demonstrated that the proteins encoded by the *uvr* genes of Arthrospira and Lactococcus lactis (see Figures 2–6) share significant similarities and homologies with ORF252 as well as with another protein of paramount importance in human DNA repair, the UV excision repair protein RAD23 homolog B, that is encoded by the gene RAD23B. This is a multi-ubiquitin chain receptor involved in modulation of proteasomal degradation. The region of interest is between residues 237 and 409 and the tri-dimensional molecular structure of this region is shown in Figures 8A and 8B.

**Fig. 8A.**
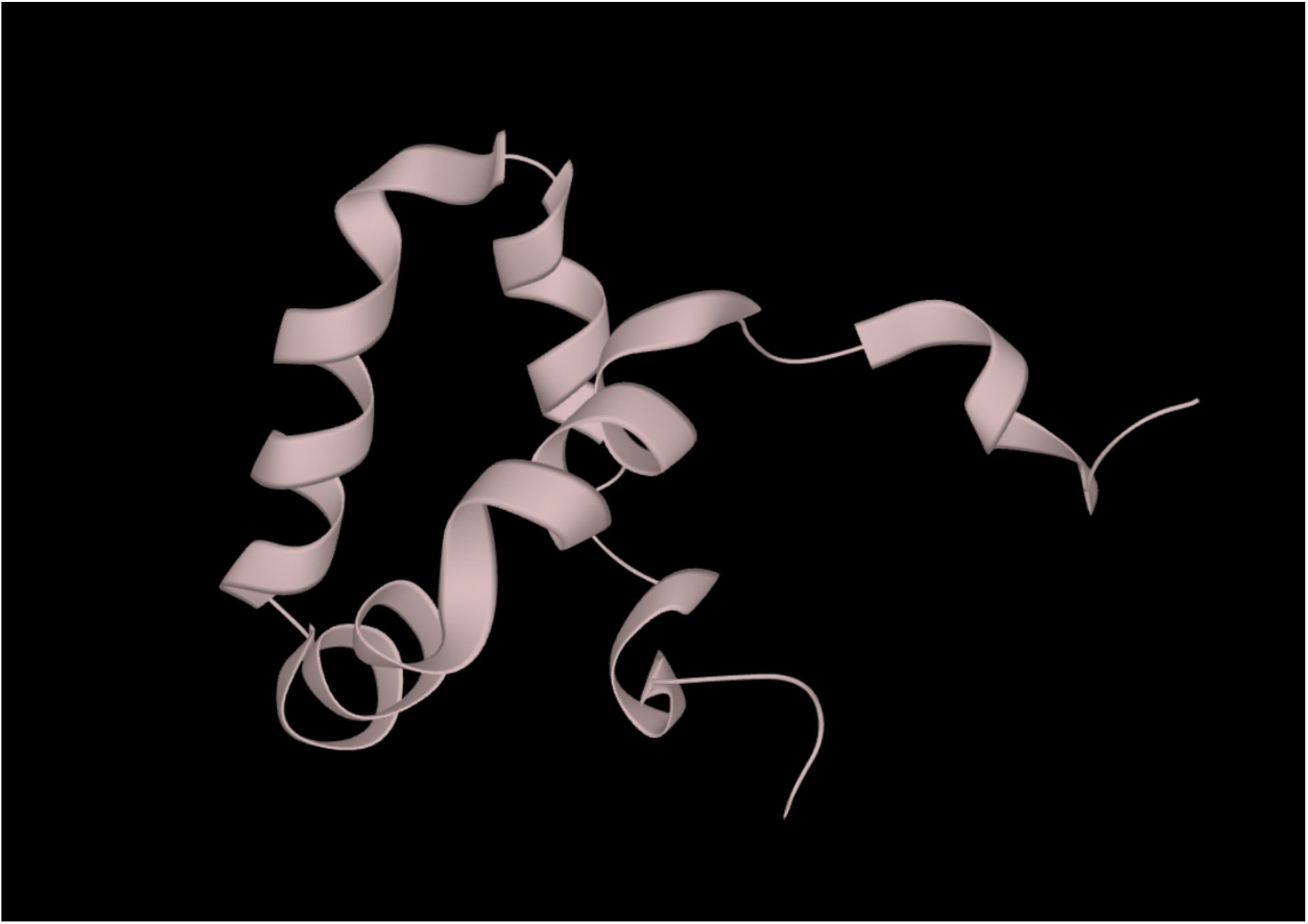
Tridimensional structure of human RAD23B, chain A, positions 275-342.

**Fig. 8B.**
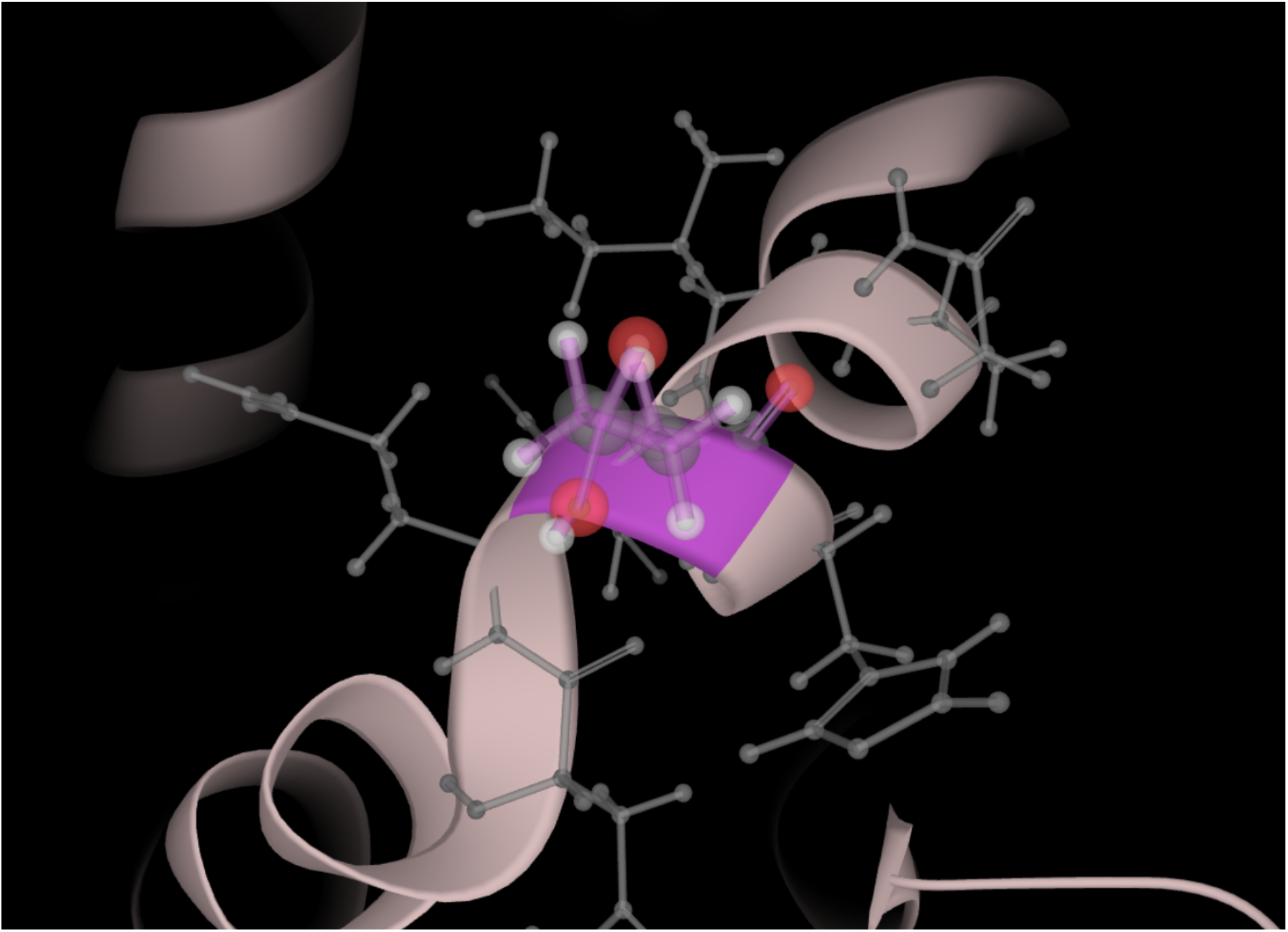
Tridimensional structure of human RAD23B, chain A, positions 275-342; magnification.

Figs. 9 and 10, show the regions where the homologies between the four proteins (UvrABC system protein B of Arthrospira platensis, UvrABC system protein B of Lactococcus lactis subsp. lactis, human RAD23, and viral ORF252) are particularly evident as highlighted by the similarity of the amino acids, the hydrophobicity profiles (Fig. 9), and the presence of polar residues (Fig. 10).

**Fig. 9.**
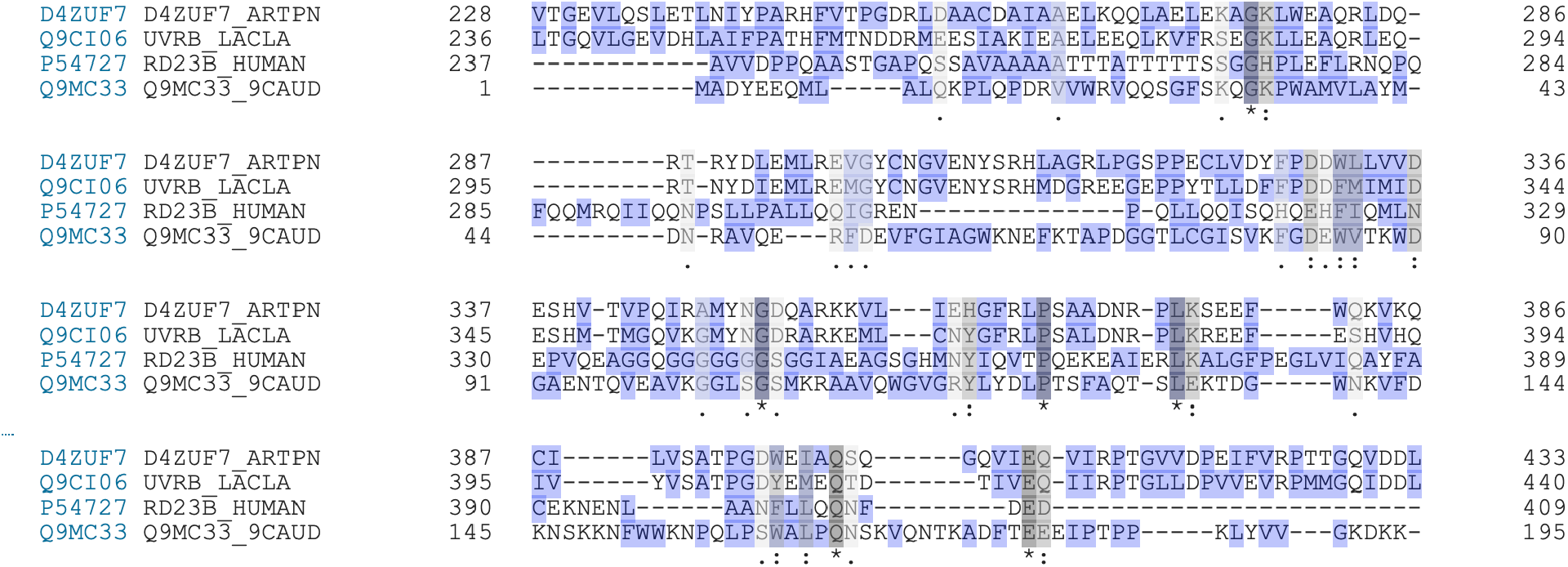
Results of alignment between UvrABC system protein B (uvrB gene) of Arthrospira platensis (first line), UvrABC system protein B (uvrB gene) of Lactococcus lactis subsp. lactis (second line), human RAD23 (third line), and Lactococcus phage ul36 ORF252 (fourth line). Residues showing significant degree of similarity are highlighted in grey. Hydrophobic residues are highlighted in indigo.

**Fig. 10.**
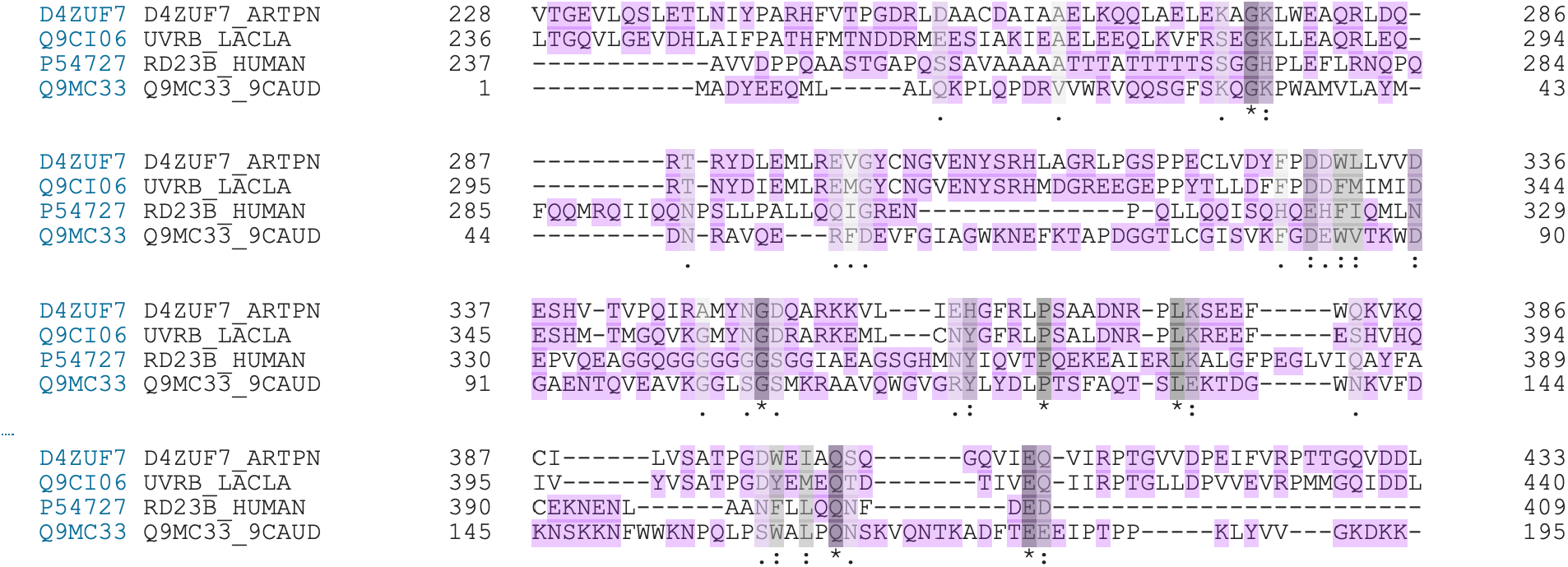
Results of alignment between UvrABC system protein B (uvrB gene) of Arthrospira platensis (first line), UvrABC system protein B (uvrB gene) of Lactococcus lactis subsp. lactis (second line), human RAD23 (third line), and Lactococcus phage ul36 ORF252 (fourth line). Residues showing significant degree of similarity are highlighted in grey. Polar residues are highlighted in lilac.

The high degree of similarity, hydrophobicity, and presence of polar residues in these particular regions of the four proteins is deemed responsible for the transfer of information between bacteria, phages and human cells. This is particularly evident when comparing the human and the viral protein (Fig. 11) where the sequence of hydrophobic and polar residues shows a very significant degree of homology.

**Fig. 11.**
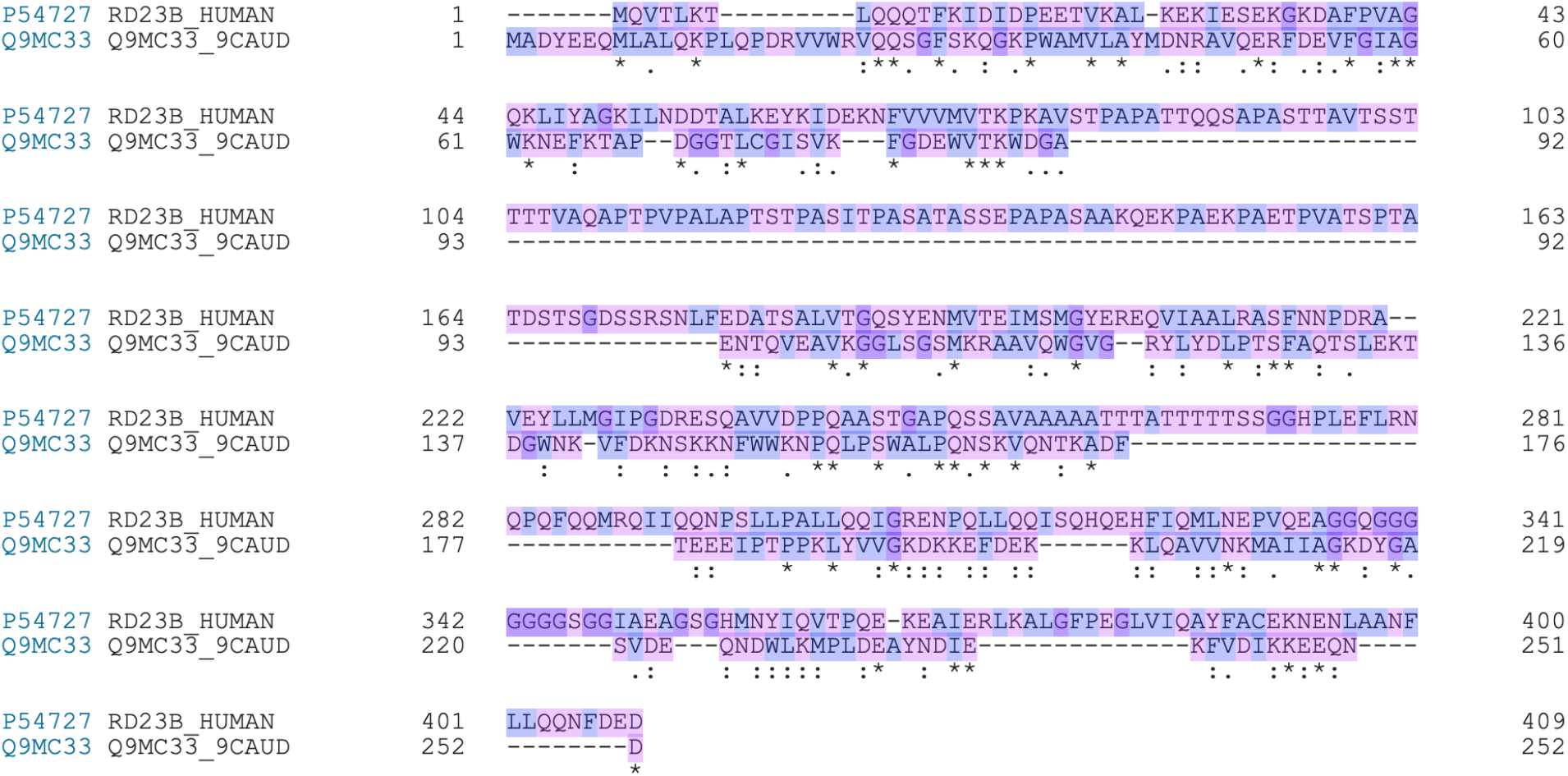
Results of alignment between human RAD23 (upper line) and Lactococcus phage ul36 ORF252 (lower line). Hydrophobic residues are highlighted in indigo. Polar residues are highlighted in lilac.

## Discussion

The results presented in this article lay the foundation for the development of a novel approach to protection against the harmful effects of ionizing and non-ionizing radiations. In addition to Arthrospira platensis, it is worth discussing the use of Equisetum arvense (field horsetail) in the co-culture with the probiotic blend described in references 30-35. This is a perennial plant from the family of Equisetaceae. Flavonoids, phenolic acids, alkaloids, phytosterols, tannins, and triterpenoids are among the most widely known phytochemical compounds of Equisetum arvense. Several articles have described different biological effects of Equisetum arvense such as anti-oxidant, anti-inflammatory, anti-bacterial, anti-fungal, blood vessel relaxant, neuro- and cardio-protective, and anti-proliferative properties (72). The anti-cancer properties of Equisetum arvense are so well documented that a recent article published in 2020 states *“[Equisetum arvense] may serve as an alternative anticancer agent for the treatment of pancreatic carcinoma with no or least side effects to the patient*” (73). Also the anti-inflammatory properties of the plant are well documented and the mechanism of action for the anti-inflammatory effect is based on modulation of interleukins and interferon (74). The rationale for using Equisetum arvense in the co-culture described above is based on two properties of the plant. 1. Its anti-oxidant properties that counteract oxidative stress caused by electromagnetic radiations, thus synergizing with Arthrospira platensis and with phages of the probiotic blend resulting in inhibition of excessive reactive oxygen species formation. 2. The high content of silica that is arranged in surface fractals in the stem and leaf of Equisetum. Silica from Equisetum arvense plays a fundamental role in the co-culture as it favors horizontal gene transfer between radio-resistant Arthrospira platensis and the microorganisms of the probiotic blend. A number of studies have demonstrated that adsorption to minerals such as silica and silicates increases DNA longevity and stability in the environment. Such DNA-mineral associations can play the role of pools of genes that can be stored across time, an essential feature in the context of an approach designed to withstand the noxious effects of electromagnetic radiations. More importantly for the transfer of biological information is the fact that DNA associated with silica is available for incorporation into organisms different from those where DNA originated through the process of horizontal gene transfer. The complex silica-DNA holds the potential for successfully transferring genetic material across environments and timescales to distant organisms up to the point that it has been hypothesizes that this process has significantly influenced the evolution of life on Earth (75). Interestingly, Equisetum is considered a “living fossil”, since it is the only living genus of the entire subclass Equisetidae, a class that was widely represented in the late Paleozoic forests and survived the largest extinction event in the history of Earth, the Permian-Triassic extinction event. In the context of the co-culture, Equisetum arvense serves the purpose of reproducing the conditions that are believed to be present at the beginning of the life on Earth, when nucleic acids bound to minerals such as silica and silicates, and were thus able to maintain, transfer, and propagate the primeval biological information. More precisely, silica favors exchange of information between radio-resistant Arthrospira platensis and the microorganisms of the probiotic blend. Such an optimization of the exchange of biological information mediated by silica occurs at the classical biochemical level through horizontal gene transfer as well as at the quantum level. It has been demonstrated that DNA self-assembles in network structures on silica surfaces in such a way that it can be used for quantum computation (76), and it has also been demonstrated that quantum entanglement synchronizes DNA strand breakage across relatively large distances by enzymes that recognize specific sequences. Such a quantum entanglement, that in the case of the co-culture is amplified by silica, explains the observed persistent effects of enzymes that interact with DNA across longer spatial separations, a phenomenon that cannot be explained in terms of classical biochemistry (77). In addition, silica serves the purpose of absorbing phages, thereby enhancing their effectiveness in transferring the information to other microorganisms as well as to human cells. It has been demonstrated that phages are active while adsorbed on the surface of silica particles and the number of active phages bound to the silica is enhanced by the presence of ionic surfaces, with greater surface charge correlating with greater concentration of adsorbed phages (78). In the co-culture, the ionic surface are represented by the phenolic acids of Equisetum arvense and the simultaneous presence of silica and phenolic acids leads to synergism as far as phage absorption is concerned.

## Acknowledgements

The Author wishes to thank the Colleagues at Silver Spring Sagl, Switzerland, for their continuous support and encouragement and for sharing scientific information.

## Ethics

This article is original and contains material that has not been submitted or published in any scientific journal. However, the concepts and some of the results presented in this article are part of a patent application to the United States Patent and Trademark Office filed in June 2020.

